# MS-PyCloud: An open-source, cloud computing-based pipeline for LC-MS/MS data analysis

**DOI:** 10.1101/320887

**Authors:** Li Chen, Bai Zhang, Michael Schnaubelt, Punit Shah, Paul Aiyetan, Daniel Chan, Hui Zhang, Zhen Zhang

## Abstract

Rapid development and wide adoption of mass spectrometry-based proteomics technologies have empowered scientists to study proteins and their modifications in complex samples on a large scale. This progress has also created unprecedented challenges for individual labs to store, manage and analyze proteomics data, both in the cost for proprietary software and high-performance computing, and the long processing time that discourages on-the-fly changes of data processing settings required in explorative and discovery analysis. We developed an open-source, cloud computing-based pipeline, MS-PyCloud, with graphical user interface (GUI) support, for LC-MS/MS data analysis. The major components of this pipeline include data file integrity validation, MS/MS database search for spectral assignment, false discovery rate estimation, protein inference, determination of protein post-translation modifications, and quantitation of specific (modified) peptides and proteins. To ensure the transparency and reproducibility of data analysis, MS-PyCloud includes open source software tools with comprehensive testing and versioning for spectrum assignments. Leveraging public cloud computing infrastructure via Amazon Web Services (AWS), MS-PyCloud scales seamlessly based on analysis demand to achieve fast and efficient performance. Application of the pipeline to the analysis of large-scale iTRAQ/TMT LC-MS/MS data sets demonstrated the effectiveness and high performance of MS-PyCloud. The software can be downloaded at: https://bitbucket.org/mschnau/ms-pycloud/downloads/

---

Rapid development and wide adoption of mass spectrometry-based proteomics technologies have empowered scientists to study proteins and their modifications in complex samples on a large scale^1,2^. However, this progress created unprecedented challenges for researchers to store, manage, and analyze the large scale mass spectrometry data. For instance, the Clinical Proteomic Tumor Analysis Consortium (CPTAC) has generated around 1.6 terabyte (TB, 240bytes) raw mass spectrometry data for TCGA ovarian cancer proteome using tandem mass spectrometry with isobaric tag for relative and absolute quantitation (iTRAQ) multiplex labelling method ^3,4^ Mass spectrometry data analysis involves high-computing demands for peptide identification through database search, protein inference, post-translational modifications (PTMs) identification, and quantification of PTMs and global proteins on large scale high throughput data. Currently, proprietary software packages such as Proteome Discovery (Thermo Scientific), Sorcerer (Sage-N Research), SCAFFOLD (Proteome Software) and PEAKS (Bioinformatics Solutions Inc.) rely on high-performance computing hardware and computer cluster management. However, their maintenance and access are typically expensive. Open source, cloud-based computing has shown its advantages in terms of scalability and financial cost ^5^. Many individual open-source tools with different search engines are available, e.g. OpenMS/TOPP ^6^, TPP ^7^, CPAS (Labkey Software Foundation) and MASPECTRAS ^8^. However, it requires additional efforts to migrate the existing pipelines to a cloud-based computing environment.

In order to integrate the various open-source tools and resources into a comprehensive yet easy to use pipeline for LC-MS/MS data processing, we developed a cloud-computing based open-source software package, MS-PyCloud, with all-inclusive customizable components: data file integrity validation, data quality control, false discovery rate estimation, protein inference, PTM identifications, and quantitation of PTMs and global proteins. A graphical user interface (GUI) is also developed for end users for easy accessibility and parameter settings. Currently, MS-PyCloud supports cloud computing on Amazon Web Services. A key advantage of MS-PyCloud is its ability to scale seamlessly for datasets of varying sizes to maintain a relatively constant turnaround time which is often needed in explorative analysis for making on-the-fly modifications of data processing parameters. MS-PyCloud was compared using data from breast cancer xenograft tissues with other proteomic pipelines to demonstrate its effectiveness. With cloud-based computing, MS-PyCloud was also able to complete within five hours the processing of a large LC-MS/MS dataset which included 44 iTRAQ sets, each with 25 LC-MS/MS runs on an Orbitrap Velos Pro mass spectrometer (1100 raw files in total, nearly 850 GB of data). The protein identification and quantitation results showed high performance and good reproducibility. The software is available for download at: https://bitbucket.org/mschnau/ms-pycloud/downloads/

## METHODS

**Schematic workflow for MS-PyCloud.** The schematic workflow of MS-PyCloud is shown in Figure 1, which includes three major components: peptide identification through database search, protein inference, and protein/PTM quantitation. After converting raw LC-MS/MS data to mzML format, the files then searched against either a standard protein database or customized database for peptide identification, using one of the publicly available search engines such as MyriMatch ^9^, MS-GF+ ^10^ and Comet ^11^. The identified peptides are represented in the format of pepXML, mzid, or XML. Peptide spectrum matches (PSMs) are filtered based on a custom-defined FDR cutoff and then the false discovery rate (FDR) is estimated for identified peptides using a decoy search. Significant PSMs from all files are grouped to infer the represented proteins parsimoniously using a bipartite graph analysis algorithm adopted in many protein inference tools^13–15^. Proteins/PTM can be quantified for labeled tags such as iTRAQ and TMT. For labeled data such as 4-plex iTRAQ, we implement a function to extract iTRAQ reporter ion intensity based on identified peptide and raw LC-MS/MS data. The peptide and protein quantification are calculated based on iTRAQ reporter ion intensity at PSM level and the corresponding FDR is estimated at both peptide and protein level. The relative protein abundance is calculated as the median of relative abundances of peptides belonging to same protein, which is calculated as the median of log2 ratio of reporter ion intensities of PSMs belonging to the same peptides^3,16,17^.

**Figure 1.**
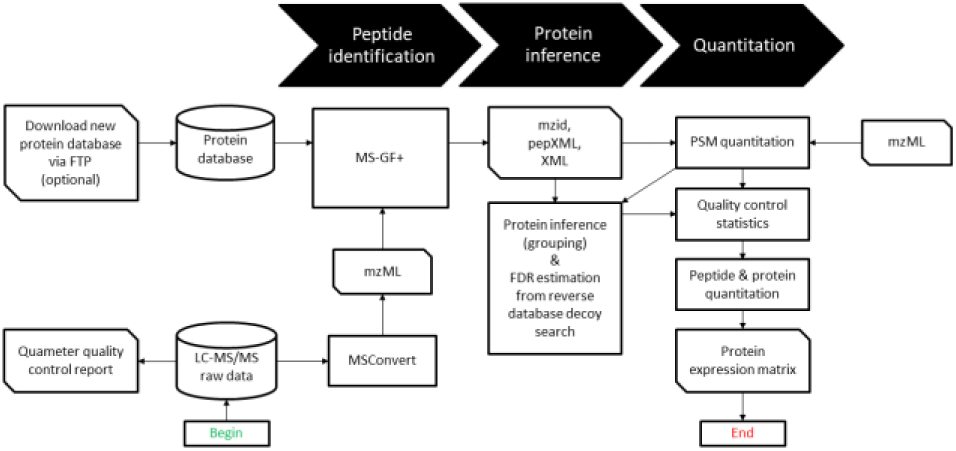
Schematic workflow of MS-PyCloud for LC-MS/MS data analysis. The orange blocks are either raw input data/database, intermediate results or final data matrix and reports. The blue blocks are open source tools/software used in the pipeline. The green blocks are the functional modules implemented in the package.

**Cloud-computing based solution.** The architecture of cloud computing of MS-PyCloud is shown in Figure 2. Amazon Web Services (AWS) is used as an example to provide scalable computing resources through Elastic Compute Cloud (EC2) and data storage using Elastic Block Storage (EBS) or Simple Storage Service (S3). StarCluster (StarCluster 2010, http://star.mit.edu/cluster/), an open source cluster-computing toolkit for Amazon’s Elastic Compute Cloud (EC2), is used for cluster management and job scheduling with the open source version of Sun Grid Engine (http://gridscheduler.sourceforge.net/index.html). On the top, the MS-PyCloud core package, composed of open-source tools, provides services of peptide identification on multiple computers in parallel. Overall, it forms a high performing, scalable, cost-effective cloud-based pipeline.

**Figure 2.**
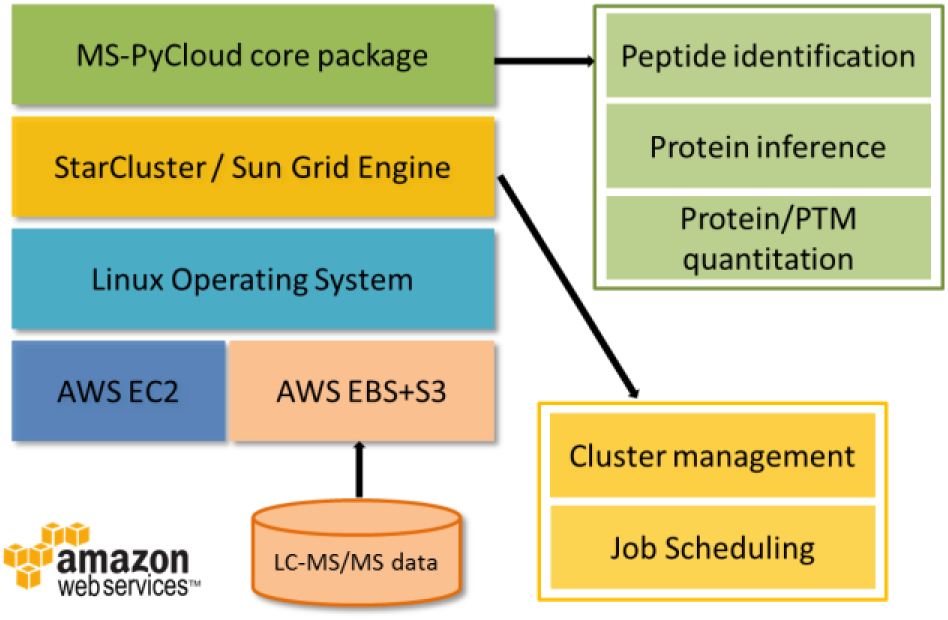
Architecture of cloud computing of MS-PyCloud pipeline based on Amazon Web Services (AWS) and StarCluster.

**Graphical User Interface.** MS-PyCloud’s graphical user interface (GUI) is also written in Python. With several user-defined configuration text files, it provides a user-friendly platform for processing mass spectrometry data. A Python program is also included that can be executed from the command line to run all pipeline steps for batch processing without using the GUI. The GUI gives the user easy access to the user-defined configuration files needed to run the pipeline, as well as information about what modifications the user has specified. The GUI is updated in real-time, as the parameters are modified in the configuration files to reflect the most recent user settings. Importantly the GUI allows the user to intuitively and swiftly set the peptide modifications present in the samples to be analyzed. In addition, the GUI allows the user to select which steps to run and whether the steps should be run on the local machine or the cloud using Amazon AWS cloud computing. The graphical interface is shown below in Figure 3.

**Figure 3.**
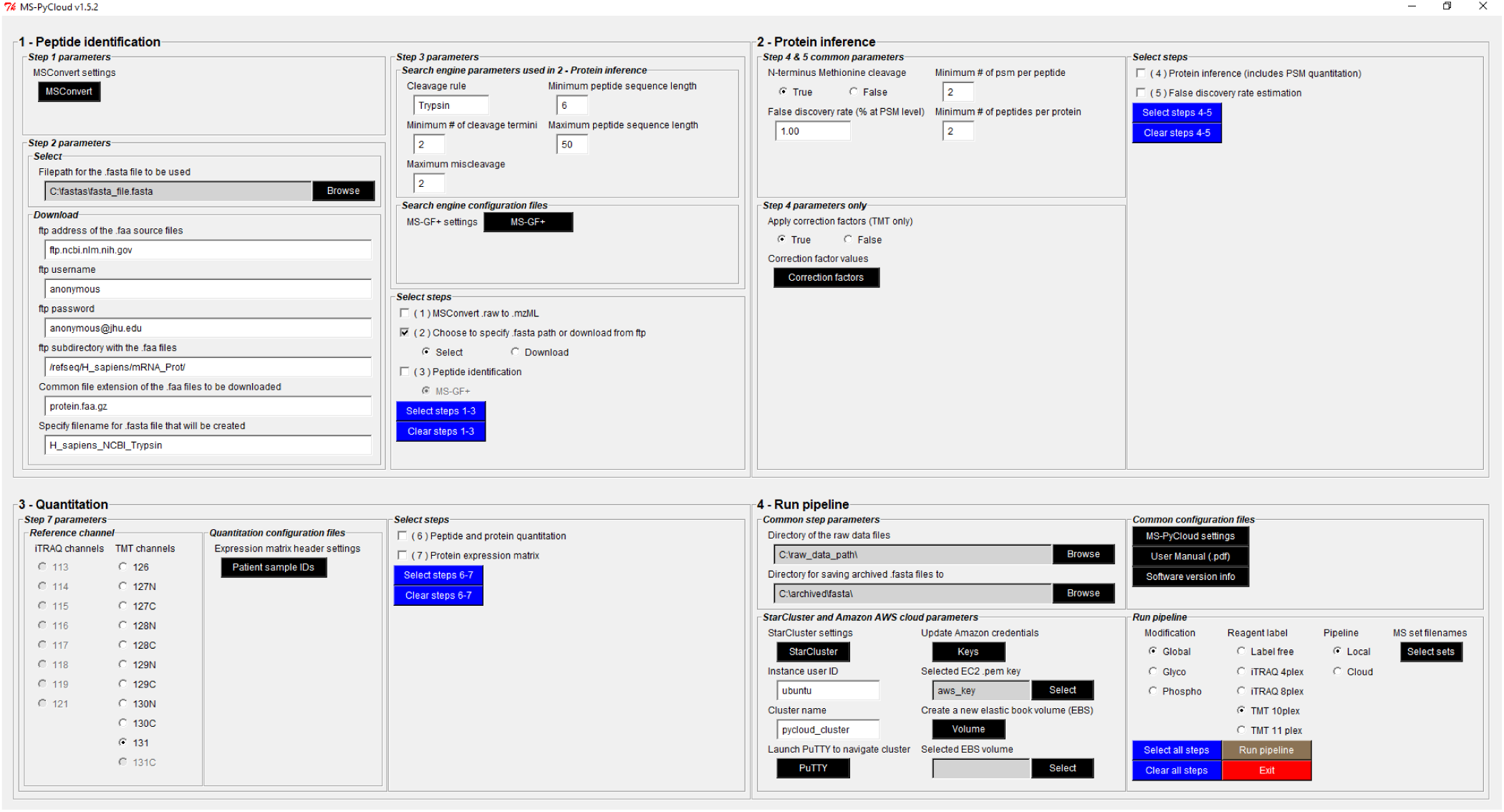
Graphical User Interface (GUI) of MS-PyCloud. Includes user-defined parameters for peptide identification, protein inference, and quantitation. Also allows the user to specify the ftp source of a fasta database to be downloaded, and define parameters for interfacing with Amazon’s Web Service through StarCluster.

**Global proteome data.** Two sets of global proteomic data were analyzed to evaluate MS-PyCloud. The first dataset was from breast cancer xenograft tissues of basal-like tumor and luminal tumor^18^. Two technical replicates of basal-like tumor were labeled by iTRAQ 114 and 115 and two technical replicates of luminal tumors were labeled by iTRAQ 116 and 117. iTRAQ labeled mixtures were fractionated 24 times by high pH reverse phase HPLC, respectively, and each fraction was analyzed by LTQ Orbitrap Velos for LC-MS/MS analysis.

The second dataset was a part of a large-scale proteomic characterization of ovarian high-grade serous carcinoma which included 122 ovarian cancer samples and 10 quality control (QC) repeats of one additional ovarian cancer sample. After protein extraction and tryptic peptide digestion, the desalted peptides were labelled with 4-plex iTRAQ reagents where channel 114 was used for labeling the pooled reference sample. A total of 44 4-plex iTRAQ experiments were run and each experiment had 24 fractions and unfractionated analysis. The global proteome data was generated by LC-MS/MS using Orbitrap Velos Pro mass spectrometer (Thermo Scientific), which includes 1,100 raw files in total, nearly 850 GB.

## RESULTS

**Analysis of breast cancer xenograft tissues using MS-PyCloud.** The pipeline was first applied on breast cancer xenograft tissues on a single workstation (Intel Xeon CPU). In the pipeline, MyriMatch ^9^ and the Human and Mouse combined protein sequence database ^18^ were used for peptide identification. Thermo RAW files were converted to mzML files by ProteoWizard msConvert. MyriMatch used 10 ppm precursor mass tolerance, considered full-tryptic peptides, allowed for isotopic error in precursor ion selection, and applied static +57 modification to cysteine, static +144 (iTRAQ-4plex) to peptide N-terminus and lysine, and dynamic +16 oxidation to methionine. The peptide spectrum matches (PSMs) identified from MyriMatch were further filtered by (1) peptide FDR less than 1%, (2) number of missed cleavage sites less than 2, and (3) at least 2 peptides for an identified protein. For comparison, we also implemented IDPicker ^13^ for protein inference based on PSMs identified from MyriMatch. In addition, a commercial pipeline, Proteome Discoverer (Thermo Scientific), was compared which used SEQUEST ^19^ for database searches. We set up similar database search parameters and PSMs filtering criteria for protein inference.

In breast cancer xenograft tissues, our pipeline identified 111,996 PSMs and 52,292 peptides, representing 6,289 unique protein groups at 1% peptide FDR. IDPicker and Proteome Discoverer (PD) identified similar numbers of PSMs, peptides and protein groups and Figure 4 shows the Venn diagram of total identification among MS-PyCloud, IDPicker and PD. The identified PSMs, peptides and protein groups from MS-PyCloud were highly overlapped with the ones identified from other two pipelines. For example, 95.95% PSMs, 97.15% peptides and 96.10% protein groups identified from MS-PyCloud were also identified either from IDPicker or PD, showing the effectiveness of our pipeline.

**Figure 4.**
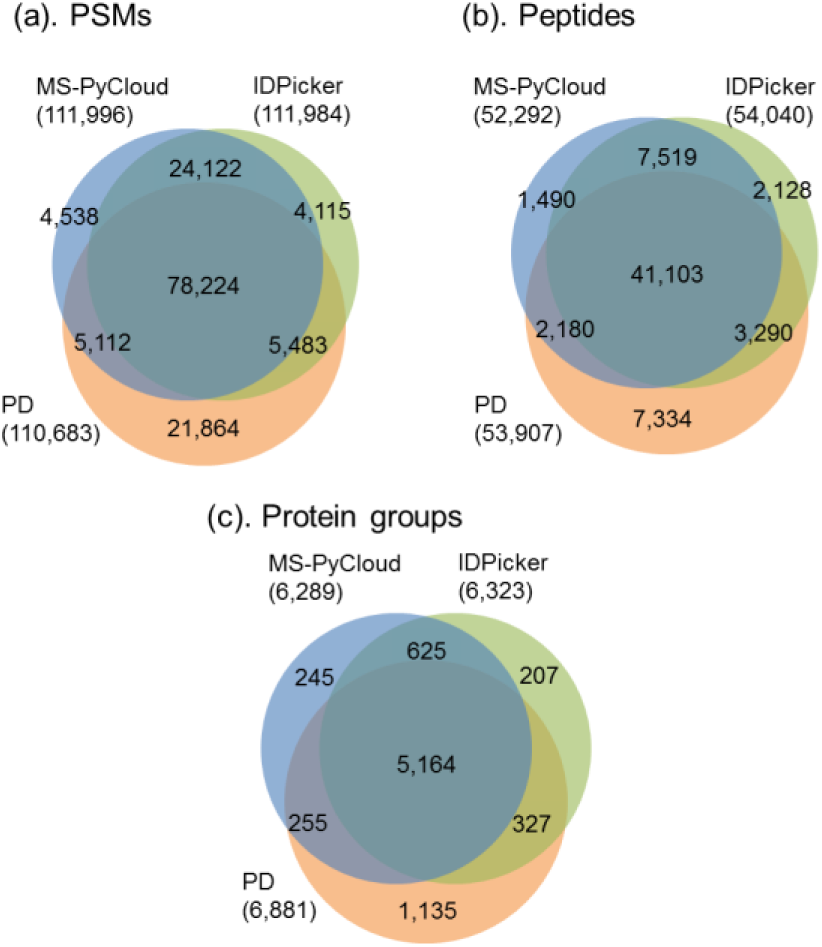
Venn diagram of total identified (a) PSMs, (b) peptides and (c) protein groups among MS-PyCloud, IDPicker and Proteome Discoverer (PD) in 24 iTRAQ runs.

Since IDPicker cannot handle the quantitation of iTRAQ labeled data, we only compared protein quantitation between MS-PyCloud and PD for the common identified protein groups. Figure 5 shows the scatter plot of relative protein abundances between MS-PyCloud and PD for two replicates and they are significantly correlated with Spearman correlation coefficients 0.94 (p_value < 0.001).

**Figure 5.**
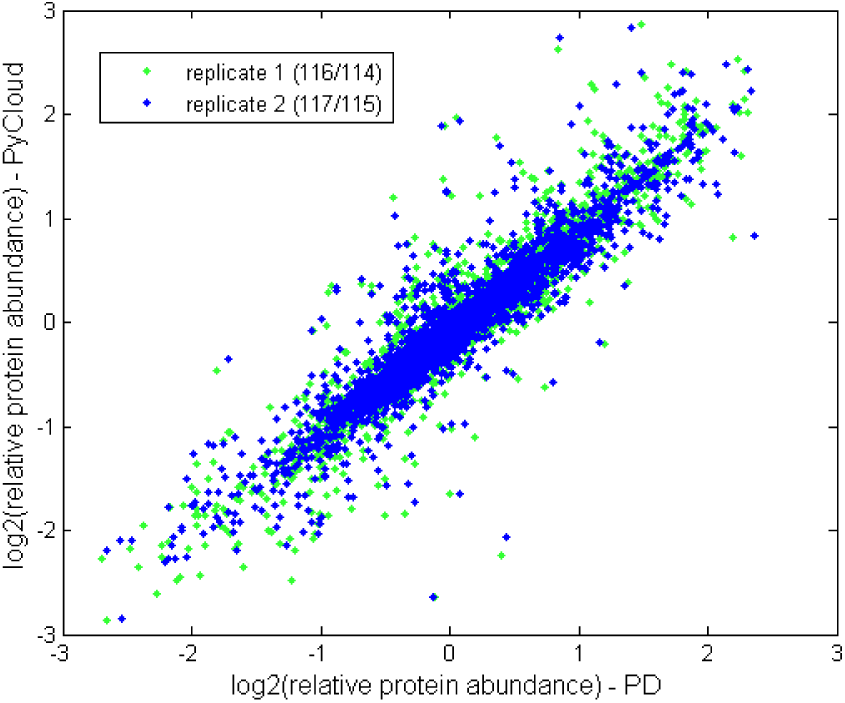
Scatter plot of relative protein abundances between basal (116, 117) and luminal (114, 115) from MS-PyCloud and PD.

**Analysis of ovarian cancer samples using cloud-based pipeline.** The cloud-based pipeline was implemented to process global proteome data of 122 ovarian cancer samples and 10 QC repeats of an additional ovarian cancer sample. In the pipeline, MyriMatch and the RefSeq Human protein sequence database (release version 54) were used for peptide identification. Thermo RAW files were converted to mzML files by ProteoWizard msConvert. The 1100 converted mzML files were uploaded to Amazon AWS EBS and 100 instances (m3.xlarge) were configured for high performance database search and protein quantitation. For global peptide search, MyriMatch used 10 ppm precursor mass tolerance, considered full-tryptic peptides, allowed for isotopic error in precursor ion selection, and applied static +57 modification to cysteine, static +144 (iTRAQ-4plex) to peptide N-terminus and lysine, and dynamic +16 oxidation to methionine. The 1100 pepxml files from MyriMatch were used for final protein assembly. The peptide spectrum matches (PSMs) were further filtered by (1) peptide FDR less than 1%, (2) number of missed cleavage sites less than 2, (3) at least 2 PSMs for an identified peptide, and (4) at least 2 peptides for an identified protein.

MS-PyCloud pipeline spent ~5 hours to generate the final protein expression matrix dataset and the total cost was around $300 for cloud computing as of 2013. The cost could be reduced 40% in 2015 according to Amazon AWS Pricing website. Global proteomics analysis of 132 ovarian cancer samples identified 4.42 × 10^6^ PSM, 122,176 unique peptides and 10,794 protein groups in total. To assess the precision of protein quantitation estimation of our pipeline, we calculated coefficient of variation (CV), defined as the standard deviation divided by mean of replicated measurements. The estimation of the CV used the relative protein abundance data from the 10 replicates of the QC sample instead of the log2 transformed data since the log2 ratio data tended to have means close to 0. Among these proteins 89.96% had a CV < 15% (Min: 0.017, 1^st^ Qu. 0.055, Median 0.075, Mean 0.090, 3^rd^ Qu. 0.101, Max. 0.950).

The short turnaround time with the use of cloud-based computing made it possible to repeatedly reanalyze such large dataset for additional features, such as to search for peptides with specific PTMs. For instance, by adding dynamic modifications on lysine residues, we were able to quickly “on-the-fly” reprocess the data and identified 3,527 unique acetylated peptides and 2,013 unique protein groups that are acetylated in the ovarian cancer samples, which provided additional data for needs prompted from initial analysis of the global protein data.

## DISCUSSION AND CONCLUSION

We have developed an open source, cloud computing-based pipeline, MS-PyCloud, for LC-MS/MS data analysis. MS-PyCloud consists of only open source software tools to ensure the transparency and reproducibility of data analysis process and results. Scalability, performance and robustness are the fundamental design considerations of this pipeline. Cloud-based solution yields very fast performance even on large data sets and the on-demand model avoids the hassles of managing private computer clusters and significantly reduces computing costs. The applications of MS-PyCloud to high throughput proteomic data have demonstrated the effectiveness and efficiency of our proposed pipeline.

MS-PyCloud shows good effectiveness in identifying peptides and protein groups from the case study on breast cancer xenograft tissues. MS-PyCloud has identified more overlapped PSMs, peptides and protein groups with IDPicker than PD since MS-PyCloud and IDPicker use the same search engine MyriMatch. The non-overlapped PSMs and peptides between MS-PyCloud and PD may be due to different search engines and different FDR calculation, and non-overlapped proteins are partially due to a different protein inference algorithm, which has been compared and studied previously^20–22^.

Cloud-computing based solution provides good features such as scalability and reliability. Recently, Trans-Proteomic Pipeline (TPP) has been enabled with Amazon Web Services cloud computing to speed up data analysis pipeline at a very low cost ^23^. An Automated Proteomics Pipeline (APP) was also implemented using Java to integrate a set of tools and allow distributed execution on most available computers with minimal setup^24^. However, existing pipelines tried to provide a uniform solution for different types of mass spectrometry data (e.g., labeled or unlabeled) generated from different instruments, which made it difficult for users to customize for their specific application. Also some pipelines assembled different tools from multiple sources, which increased the computational burden on I/O to transfer the intermediate results. MS-PyCloud implements all key components using Python, which makes it easy to incorporate existing Python packages for MS data analysis or be accessed by other developing packages.

Cloud-computing based solution has shown its advantage in speeding up data analysis cycles with reduced turnaround time that is conducive for scientists to test “what-if’ scenarios with different parameters and settings. Furthermore, with high computing capacity, it is possible to carry on *De novo* peptide identification on the mass spectrometry data and facilitate biological discovery.

## Author Contributions

All authors have given approval to the final version of the manuscript.

## Notes

The authors declare no competing financial interest.

## ACKNOWLEDGMENTS

This work was supported by National Institutes of Health, National Cancer Institute, The Clinical Proteomic Tumor Analysis Consortium (CPTAC, U24CA160036), the Early Detection Research Network (EDRN, U01CA152813).

